# ISdetector: precise mapping of insertion sequences and associated structural variations from short-read sequencing data

**DOI:** 10.64898/2026.03.18.712784

**Authors:** Yang Zhou, Bingxin Lu, Yanlin Zhao

**Affiliations:** Chinese Center for Disease Control and Prevention, No. 155 Changbai road, Changping district, Beijing, China, 102206; School of Biosciences, University of Surrey, Stag Hill, Guildford, UK, GU2 7XH

## Abstract

**Motivation:** Repetitive insertion sequences (ISs) and adjacent structural variants (SVs) complicate precise mapping from bacterial short reads. Accurate insertion coordinates are important for studying genome evolution and tracking transmission.

**Results:** ISdetector constructs IS-specific references with target copies removed and clusters soft-clipped reads to pinpoint insertion sites. On simulated genomes, it had higher recall and F1 score than ISMapper and MGEFinder. On 19 *Mycobacterium tuberculosis* genomes, it achieved the highest high-confidence F1 score (0.86). On five *Shigella sonnei* genomes, it performed comparably to ISMapper (F1 score 0.85). It detected 81 of 105 IS6110 copies with associated SVs or complex events, versus 41 (ISMapper) and 12 (MGEFinder). With four threads, it outpaced both comparators at any thread count.

## 1 Introduction

Insertion sequences (ISs) are the smallest and simplest mobile genetic elements (MGEs) found abundantly in prokaryotic genomes. They typically range from hundreds to a few thousand base pairs in length and only carry the genetic information necessary for their own mobility (Siguier, Filée and Chandler, 2006). They are capable of repeated insertion into many different sites within a genome using mechanisms that do not require large regions of DNA sequence homology between the IS and the target site (Siguier *et al*., 2015).

As drivers of genomic plasticity, IS elements play a fundamental role in bacterial evolution by mediating diverse genetic rearrangements, including gene inactivation, inversions, and large-scale deletions (Durrant *et al*., 2020; Kirsch, Hryckowian and Duerkop, 2024; Ngan *et al*., 2024; Zhou *et al*., 2025). Their transposition can modulate gene expression and bacterial fitness. For example, IS6110 insertions upstream of the *phoP* virulence gene or within the *plcABC* phospholipase operon actively modulate host-pathogen interactions and macrophage survival in *Mycobacterium tuberculosis* (MTB) (Hawkey *et al*., 2015; Farzand *et al*., 2022). In addition, the accumulation of ISs has influenced many aspects of genome evolution in intracellular pathogenic Shigella species (Seferbekova *et al*., 2021). IS elements also serve as critical markers for molecular epidemiology in tracking transmission chains and monitoring outbreaks in pathogens (Peres *et al*., 2018; Shitikov *et al*., 2019). Therefore, precisely mapping ISs at the population level is critical for understanding the whole picture of genetic diversity, as the exact genomic location of an IS element is a profound driver of bacterial adaptation, virulence, and outbreak dynamics.

The advent of whole-genome sequencing (WGS) has revolutionized bacterial genotyping, offering a comprehensive alternative to traditional methods. WGS data allows for the simultaneous analysis of single nucleotide polymorphisms, drug resistance profiles, and core genome phylogenies (Meehan *et al*., 2019). However, the automated detection of IS elements from high-throughput short-read data remains a significant computational challenge. IS elements are repetitive by nature, and short sequencing reads (e.g., Illumina) derived from these regions often map to multiple genomic locations, confounding standard alignment algorithms and leading to fragmented assemblies. Moreover, IS elements frequently generate adjacent deletions and inversions during the formation and resolution of cointegrates, creating structural complexities that inherently limit the resolution of general-purpose structural variation (SV) callers (Roychowdhury, Mandal and Bhattacharya, 2015; Seferbekova *et al*., 2021).

Current bioinformatics tools for IS detection each have distinct limitations. Assembly-based tools or web services, such as ISfinder and ISEScan, rely on complete or high-quality draft genomes for IS annotation (Siguier *et al*., 2006; Xie and Tang, 2017). However, short-read assemblers frequently collapse repetitive IS regions or place them on degenerate contigs, leading to incomplete or inaccurate annotations. In contrast, mapping-based tools, such as ISMapper and MGEFinder, locate insertion sites relative to a reference genome, but often struggle to resolve complex structural variations (SVs) or insertions within repetitive regions (Hawkey *et al*., 2015; Durrant *et al*., 2020). Recently, DeepMobilome used a convolutional neural network to achieve higher F1 scores than ISMapper and MGEFinder for identifying target MGEs in microbiome data (Cho *et al*., 2025), but it reports only presence or absence and not the exact insertion sites of ISs. While IS-specific tools struggle to resolve complex SVs, general-purpose SV callers, such as DELLY and LUMPY, also find medium-to-large insertions challenging and often fail to distinguish IS-mediated events from other genomic rearrangements (Rausch *et al*., 2012; Layer *et al*., 2014) . No existing tool bridges these two approaches to identify IS-mediated SVs from short reads. Together, these limitations highlight the need for specialized pipelines capable of leveraging raw short-read data to accurately map IS insertion sites and their associated SVs without relying on prior genome assembly.

To address these challenges, we present ISdetector, a comprehensive pipeline designed to identify insertions of specific ISs and associated SVs directly from paired-end or single-end sequencing data. ISdetector overcomes the limitations of ambiguous short-read alignment by constructing IS-specific clean references and achieves precise detection of IS by centroid-based identification. ISdetector enables the rapid and accurate profiling of the IS variations with associated SVs at the population level, reinstating the high discriminatory power of IS-based typing in the genomics era.

## 2 Methods

### 2.1 Overview of ISdetector workflow

We designed ISdetector (v1.1) as an automated bioinformatics pipeline to detect known ISs and associated SVs from high-throughput short-read sequencing data. We also optimized ISdetector for computational efficiency so that it can be run at the population level.

ISdetector takes as input raw sequencing reads (paired-end or single-end FASTQ), a database of query IS sequences (FASTA), and a reference genome (GenBank or FASTA). The workflow proceeds through four distinct stages (**Figure 1**). The main novelty of ISdetector lies in its two central stages: generating synthetic IS free reference sequences and clustering IS relevant reads from the mappings, which together enable more accurate localization of ISs and associated SVs.

**Figure 1.**
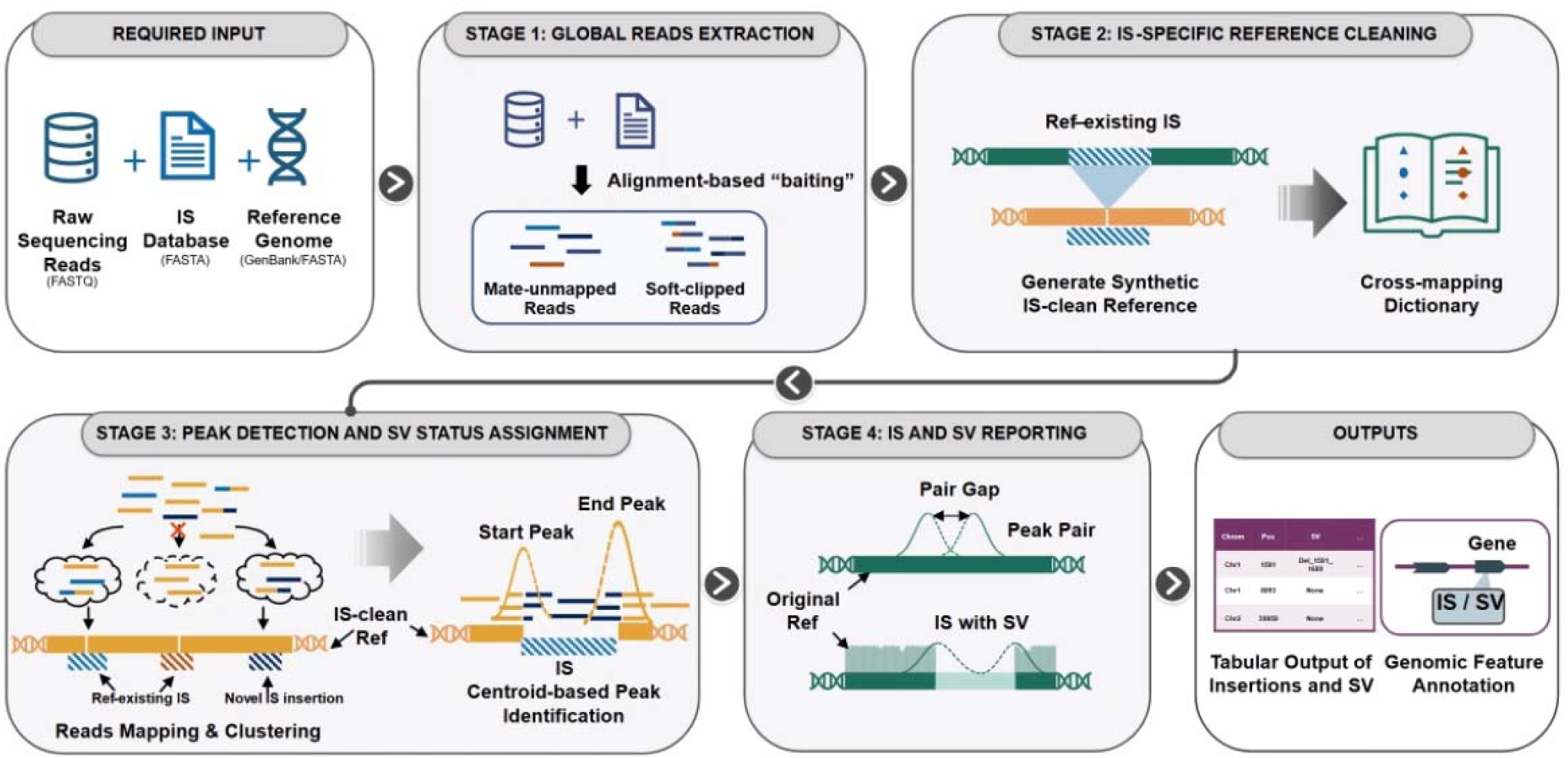
Workflow of ISdetector. ISdetector requires three input files: the raw sequencing reads, the IS database file, and the reference genome. **Stage 1**: GenBank files are automatically converted to FASTA format, and feature annotations are extracted for final reporting. Two types of reads are extracted: (1) soft-clipped reads and their mates, and (2) read pairs in which one mate maps to an IS and the other is unmapped. **Stage 2**: for each IS, the specific sequence is aligned to the reference genome and the identified IS regions are removed to generate an IS-specific clean reference. A dictionary records coordinate shifts to lift over detected positions from the synthetic clean reference back to the original genome coordinates. **Stage 3**: extracted reads are mapped to the clean reference and clustered; soft-clipped segments are locally aligned to the IS to determine precise IS coordinates, and peaks are defined by the centroid of insertion signals. Deviations in the left-to-right flanking depth ratio are used to flag adjacent deletions. **Stage 4**: peak coordinates are lifted back to the original reference, and peaks are paired using a dynamic gap threshold to determine whether they represent the two ends of a single insertion event. Insertions and associated SVs are reported in tabular form, with affected genes annotated when a GenBank reference is supplied. Alt text: Workflow diagram showing the inputs, read processing and mapping stages, and reported insertion sites.

#### Stage 1: Global read extraction

ISdetector performs a “baiting” step to extract only reads relevant to the IS elements. Input reads are aligned to the global IS database using BWA-MEM. The pipeline parses the resulting alignment to retain read pairs containing soft-clipped segments (MIN_ANCHOR_LEN=20 bp) indicative of IS unctions, or unmapped reads whose mates successfully mapped to an IS sequence.

#### Stage 2: IS-specific reference cleaning

For each IS element in the database, the pipeline generates a specific “clean” reference genome to facilitate accurate mapping of flanking regions. The specific IS sequence is aligned to the original reference genome using blastn with high stringency thresholds (default >95% identity and >90% alignment coverage). Identified IS regions within the reference are removed to create a synthetic “clean” genome. A cross-mapping dictionary is generated to record coordinate shifts by subtracting the length of removed IS elements, allowing the pipeline to “lift over” detected positions from the clean reference back to the original genomic coordinates.

#### Stage 3: Peak detection and SV status assignment

Extracted reads from Stage 1 are mapped to the IS-specific clean reference. The pipeline groups reads into clusters based on genomic proximity (CLUSTER_DIST = 50 bp and MIN_CLUSTER_SUPPORT = 20 reads) and paired reads. For each cluster, “Insertion Signals” are generated by analyzing soft-clipped segments; these segments are locally aligned to the IS query to determine the precise IS coordinate and strand orientation. Centroids for these clusters are calculated from the median genomic position and the IS coordinate. We refer to these centroids as peaks, representing the inferred IS insertion sites. A peak is called when a cluster has at least five supporting soft-clipped reads (MIN_PEAK_SUPPORT = 5), each of which must: (i) carry anchors of ≥20 bp on both the IS and the genomic flank; (ii) align to the IS with ≥90% clip coverage; (iii) have a local alignment score of at least 1.5 × the clipped length; and (iv) agree in genomic position and IS coordinate within 30 bp (PEAK_DISTANCE = 30). ISdetector also screens each peak for adjacent deletions using flanking read depth. A deletion is called if the left-to-right flanking depth ratio deviates from its expected value by more than DEPTH_DIFF_THRESHOLD (0.3), or if the clip count falls below ZERO_THRESH (default 3).

#### Stage 4: IS and SV reporting

In the final stage, peak coordinates are mapped back to the original reference genome using the cross-mapping dictionary generated at Stage 2. Specifically, the coordinates are lifted by adding back the lengths of all removed ISs located upstream of the peak. If the peak lies within ±5 bp of a reference IS, it is classified as a known reference insertion. Otherwise, it is classified as a novel insertion. To resolve full insertion events, two peaks are paired into a single call when they have the same orientation, mark opposite IS ends (Start/End), share no supporting reads, and lie within a distance threshold. For known insertions in the reference, we set the threshold to the length of IS sequences. For novel insertions, the threshold is controlled by parameter PAIR_GAP (default 20 bp), which is used to determine whether the two peaks represent the two ends of a single insertion event. For peaks assigned SV status at stage 3, their orientation, pairing status and distance (MAX_SV_SIZE =30,000) are investigated to construct final reported SVs. Unpaired peaks are still reported as low-confidence calls and marked with an asterisk. Final outputs, including insertion position, orientation, gap size, and associated genomic features, are exported to tabular reports. If a GenBank file is provided as the reference genome, the genes affected by each insertion are annotated. All genes impacted by deletions are also reported.

### 2.2 Other tools used for evaluation

We reviewed recently published IS detection tools (**Supplementary Table ST1**) and selected those that (i) take FASTQ or BAM files as input without requiring *de novo* assembly; (ii) report the exact position of every inserted copy rather than presence/absence only; and, ideally, (iii) report IS copies already present in the reference genome. We chose ISMapper (v2.0) and MGEFinder (v1.0.6) for comparisons with ISdetector. ISMapper is designed to identify the precise site and orientation of IS insertions directly from paired-end short-read sequencing data (Hawkey *et al*., 2015). MGEFinder is a bioinformatic toolbox designed to identify a wide variety of integrative MGEs and their insertion sites from short-read sequencing data of bacterial isolates (Durrant *et al*., 2020). It has two modes: the *de novo* workflow mode and the database-only mode. As the purpose of developing ISdetector is to identify targeted ISs, we only evaluated MGEFinder in database-only mode.

### 2.3 Datasets

We evaluated ISdetector and the other tools on both simulated and real genomes.

#### 2.3.1 Simulated genomes

For simulated genomes, we chose six reference genomes of different species (**Table 1**) and 15 ISs from these genomes for simulation. We obtained the representative IS sequences from the ISfinder database and aligned these sequences to each genome using blastn (v2.9.0+) (Chen *et al*., 2015) with an e-value threshold of 1e-5 to obtain the complete profile and exact insertion sites for all chosen ISs in all reference genomes.

**Table 1.**
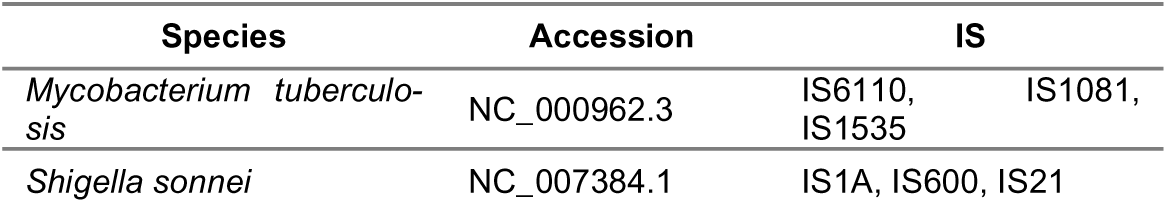
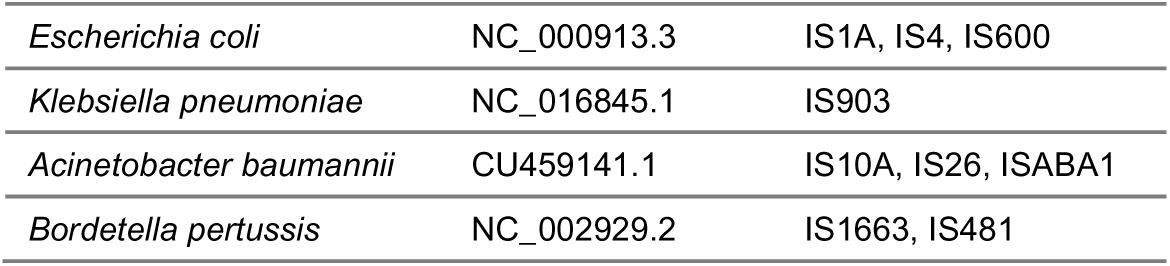
Reference genomes for IS insertion simulation.

The simulation was done using the in-house script *random_genome_variator.py*, which can be downloaded from the online git repository. Refer to the supplementary material *random_genome_variator_template.json* for the detailed parameters used for simulation. To evaluate the performance of each tool in different scenarios, we deliberately increased the weight of rare IS insertion event types, such as consecutive insertions, nested insertions, adjacent IS of different types, and IS insertions with deletions, insertions and inversions in simulated genomes. For the full list of insertion event types, refer to **Supplementary Table ST2**.

We created five replicates for each reference genome. Of these five replicates, two replicates were used in depth sensitivity analysis and parameter optimization; the other three replicates were used in batch evaluation. The script produces a TSV file for each simulated genome with details of coordinates and IS types, which serve as the ground truth database for simulated genomes in addition to the reference genome annotation GenBank files.

Paired-end reads of 150 bp were simulated using wgsim (v1.10) (Li, 2026b) at 150× depth for each simulated genome. For depth analysis, we used seqtk (v1.3-r106) (Li, 2026a) to down-sample simulated short reads to an average depth of 20×, 30×, 50×, and 100× for two replicate genomes of each reference genome. ISdetector was run using a default initial parameter profile for depth analysis.

#### 2.3.2 Real genomes

For real genome evaluation, we chose *Shigella sonnei* (*S. sonnei*) and MTB as the benchmarking species for evaluation. *S. sonnei* shows massive IS infestation (200–600 copies per genome), and MTB is notorious for its high GC content (∼65%), which are ideal candidates for evaluating robustness and efficiency.

We downloaded five complete genomes of *S. sonnei* from NCBI (NZ_CP026802.1, NZ_CP045526.1, NZ_CP049175.1, NZ_CP151334.1, NZ_CP151391.1) and simulated Illumina short-read sequencing data using wgsim in the same way as for simulated genomes at 50× depth. Ss046 (NC_007384.1) was used as the reference genome for *S. sonnei* for tools to detect IS insertions.

We selected 19 clinical MTB strains from the in-house strain bank and performed WGS on the Oxford Nanopore platform for long-read sequencing and Illumina HiSeq platform for short-read sequencing. Illumina short reads were down-sampled to 50× depth using seqtk as described. Autocycler (v0.4.0) (Wick, Howden and Stinear, 2025) and Polypolish (v0.6.0) (Wick and Holt, 2022) were employed to generate high-quality complete genomes for the 19 MTB samples, which were used to generate the ground truth database in the next step. H37Rv (NC_000962.3) was used as the reference genome.

For MGEFinder, the alignment file of short reads to the reference genome is required as the input. BWA-MEM (v0.7.17-r1188) (Li and Durbin, 2009) was used to generate the alignment file in BAM format and SAMtools (v1.10, using htslib 1.22.1) (Li *et al*., 2009) was used to generate the index.

### 2.4 Ground truth database of real genomes

For real genomes, we chose five IS families in *S. sonnei* for evaluation: IS1A, IS3, IS600, IS2, and IS21. For MTB, we chose IS6110, which is the most abundant and active IS, as the target for evaluation. We obtained the representative sequences and the complete profiles as described previously.

In the output of each tool, the insertion sites are reported as the coordinates in the reference genome. Therefore, the insertion sites detected by blastn in the real genomes must be lifted over to the corresponding coordinates in the reference genome. For this purpose, we first used a modified version of minimap2 (v2.24-r1122) (Li, 2018) to align each sample genome to the reference genome. Specifically, we added a local rescue mode to remove distortion of alignment by nearby IS sequences (all rescued records can be viewed at the git repository). We then lifted over the coordinates using an inline PAF-to-chain converter and UCSC tools (version 482) (Casper *et al*., 2026), including chainSort, chainPreNet, chainNet, netSyntenic, netChainSubset, chainStitchId, and chainSwap, followed by direct chain-based coordinate conversion (script can be downloaded from the git repository). If multiple locations are matched by the lift-over process, such as in the case of IS flanking inversions, a match to any one location was considered true but counted only once in the statistics.

### 2.5 Benchmarking metrics

Outputs of each tool were compared to the ground truth database. A reported record was counted as a true positive (TP) if it lay within ±50 bp of a ground-truth record; reported records with no ground-truth match were false positives (FPs); and ground-truth records with no matching report were false negatives (FNs). Recall was calculated by dividing the number of TPs by the number of ground truth records. Precision was calculated by dividing the number of TPs by the number of predictions (sum of the number of TPs and FPs). F1 score, the harmonic mean of recall and precision, was calculated as:

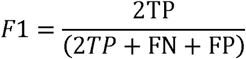

To evaluate the accuracy of each tool in predicting the insertion sites, we calculated recall/precision/F1 score for each tool with stepped thresholds of 5 bp, 10 bp, 25 bp, and 50 bp for the simulated genome dataset.

We used the real MTB genome dataset for computing resource requirements analysis. We ran each tool on Ubuntu 20.04 with Intel Xeon CPU E5-2650 v4 at 2.20 GHz with 48 cores and 252 GB of memory, varying the number of threads and recording their running time and peak RAM usage.

### 2.6 Parameter optimization

After the depth sensitivity analysis was performed, parameter optimization was run at the chosen depth across the same 12 replicate genomes used for depth sensitivity analysis.

Nine parameters of ISdetector were tested at three values each—the default plus one lower and one higher setting (**Supplementary Table ST3**). The standard sweep contained 19 ISdetector profiles: the default profile plus 18 one-factor variants which changed one parameter at a time, retaining all other default values.

Profile selection used high-confidence calls only. Each parameter’s default, lower, and upper values were ranked independently by F1 score and recall. Then we combined the nine per-parameter winners into two merged best profiles: recall best and F1 best. The merged best profiles were rerun across the same optimization panel and compared to the highest overall F1 score profile, the highest overall recall profile, and the default profile.

### 2.7 Modeling of factor impact on recall

We used ridge-penalized logistic regression to examine which factors were associated with recall for each tool. Each observation was a simulated truth event, classified as detected or missed using a 50-bp positional tolerance. We evaluated the models using repeated nested cross-validation, keeping all events from each genome replicate in the same fold. To assess the contribution of each predictor block, we refitted the model without that block and measured the increase in held-out log loss. Predictor definitions, model fitting, and evaluation are described in Supplementary Methods.

## 3 Results

### 3.1 Simulation of IS insertions

For each of the six reference genomes, the in-house script generated five replicate genomes. This produced 30 mutated FASTA files and 30 ground truth TSV files, which can be downloaded from the online git repository.

Across the 30 simulated replicate genomes, 36,495 IS copies were inserted in addition to 2,059 reference-existing ISs, falling into 13 event types with copy numbers per category ranging from 407 to 5,662 (**Supplementary Table ST4**). Additionally, 13,696 SNPs and 483 non-IS background SVs were simultaneously simulated in non-overlapping regions.

### 3.2 Batch evaluation on simulated genomes

We selected 50× sequencing depth for parameter optimization and batch evaluation because sensitivity improved most between 20× and 50×, and high-confidence calls retained 91.6% precision at 50× (**Supplementary Figure SF1**). At 50×, we selected the recall_best profile for batch evaluation. Compared with the default profile, it increased high-confidence recall from 55.1% to 59.5%, with a small decrease in F1 score from 0.69 to 0.68 (**Supplementary Figure SF2**).

The batch-evaluation panel comprised 18 simulated genomes with three replicates for each reference genome. Paired-end read datasets at 50× depth were generated from the simulated genomes. In total, the panel comprised 22,029 IS inserted copies and 987 reference-existing ISs.

#### 3.2.1 Overall performance

At 50-bp tolerance, ISdetector had higher recall and F1 score than both ISMapper and MGEFinder, but lower precision for high-confidence and all calls (**Figure 2** and **Table 2**). Overall, ISdetector provided the best balance of sensitivity and precision, whereas MGEFinder and ISMapper prioritized precision. When including confidence calls, ISMapper’s recall exceeded that of MGEFinder but remained lower than that of ISdetector.

**Figure 2.**
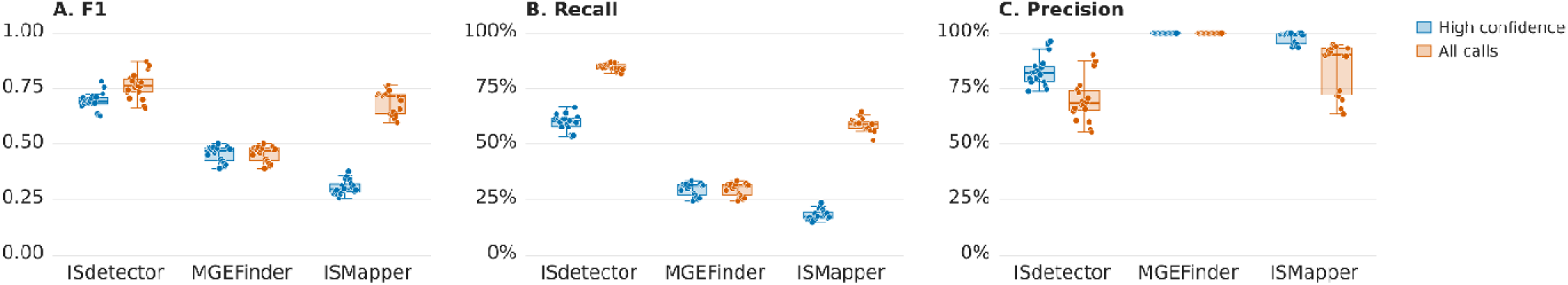
Per-genome performance of ISdetector, MGEFinder and ISMapper on simulated genomes (n=18; three replicates of each of six reference genomes, 50× paired-end reads). A. F1 score, B. Recall and C. Precision at 50-bp positional tolerance for high-confidence calls (blue) and all calls (orange). Boxes show median and interquartile range (IQR); whiskers extend to 1.5 × IQR; points are individual genomes. Alt text: Box plots compare precision, recall, and F1 scores by tool across simulated genomes.

**Table 2.** Overall performance on simulated genomes.

|  | All Confidence |  |  | High Confidence |  |  |
|---|---|---|---|---|---|---|
|  | Precision | Recall | F1 score | Precision | Recall | F1 score |
| ISMapper | 82.15% | 59.02% | 0.69 | 97.36% | 18.09% | 0.31 |
| ISdetector | 66.61% | 84.64% | 0.75 | 80.89% | 59.33% | 0.68 |
| MGEFinder | 100.00% | 29.32% | 0.45 | 100.00% | 29.32% | 0.45 |

ISdetector showed highly stable performance and achieved the highest F1 scores (0.683–0.685 for highconfidence calls and 0.744–0.746 for all calls; values below are given in the same order) across all thresholds, indicating robust breakpoint localization (**Supplementary Figure SF3**). ISMapper was substantially more tolerance-sensitive for low-confidence and imprecise calls, with marked improvements in recall (0.163-0.181, 0.374-0.590), precision (0.878-0.974, 0.520-0.821), and F1 (0.275-0.305, 0.435-0.687) up to 25 bp and only modest additional gains at 50 bp, whereas MGEFinder showed nearly constant recall (0.293) and F1 score (0.453) across all thresholds, both lower than those of ISdetector, and the highest precision (1.0) among all tools. ISdetector showed the broadest sensitivity across simulated event types and the highest recall and F1 score across the 15 IS–genome combinations. See **Supplementary Table ST5** and **Supplementary Figure SF4** for detailed results.

#### 3.2.2 Factor importance analysis

All models converged without errors and showed strong discrimination separating truth events likely to be reported from those likely to be missed (**Supplementary Table ST6**). For all tools, insertion event type produced the largest increase in log loss, suggesting that the structural form of an insertion event strongly affected recall for all three tools (**Figure 3**). However, the importance of each factor showed different orders and levels for different tools. For ISMapper, event type produced a much smaller increase in log loss than for the other two tools, and removing retained fraction increased log loss more than removing nearest-IS distance, whereas the opposite pattern was observed for ISdetector and MGEFinder; event size and IS family also showed modest log loss increase for ISMapper but none for ISdetector and MGEFinder.

**Figure 3.**
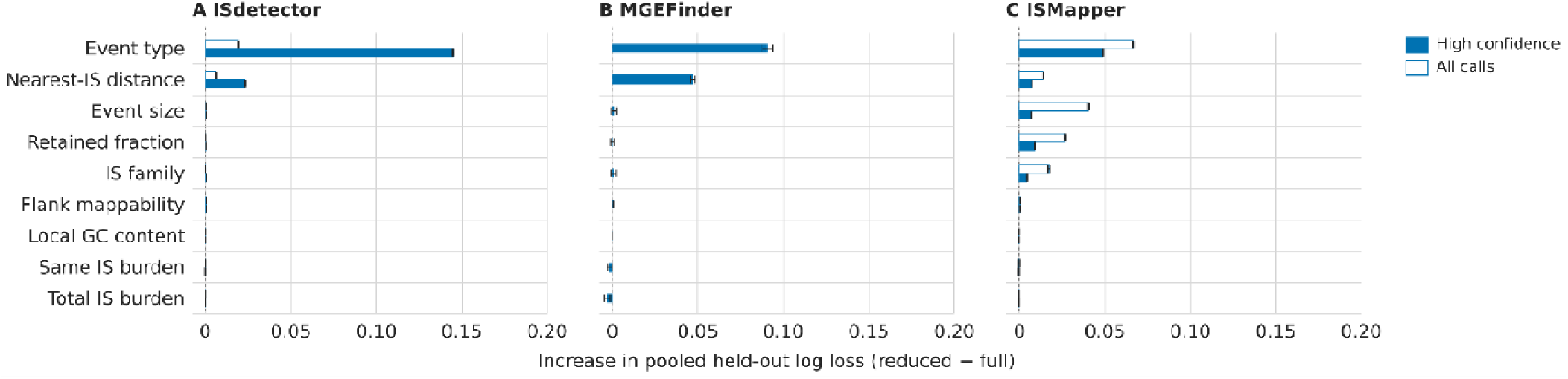
Factors associated with recall on simulated genomes. For A. ISdetector, B. MGEFinder and C. ISMapper, ridge-penalized logistic regression models predicted whether each simulated truth event was detected (50-bp tolerance). Bars show the mean increase in pooled held-out log loss when each predictor block is removed, relative to the full model, across 10 repetitions of nested cross-validation, for high- confidence calls (filled) and all calls (open). Error bars show one sample standard deviation. Alt text: Bar charts compare how much held-out log loss increases when predictor groups are removed from the models.

### 3.3 Benchmarking on real MTB and *S. sonnei* genomes

#### 3.3.1 Performance on 19 real MTB genomes with real short-read sequencing data

There were 268 copies of IS*6110* insertions in these 19 MTB genomes. ISdetector outperformed ISMapper and MGEFinder for both high-confidence and all calls (**Figure 4**). We observed more complex insertion event types in real genomes as shown in **Table 3**. “Simple insertion” was the most abundant event type in the real MTB genomes, and ISdetector showed notably superior performance to the other tools for both high-confidence and all calls, while all three tools showed poorer performance in consecutive events.

**Figure 4.**
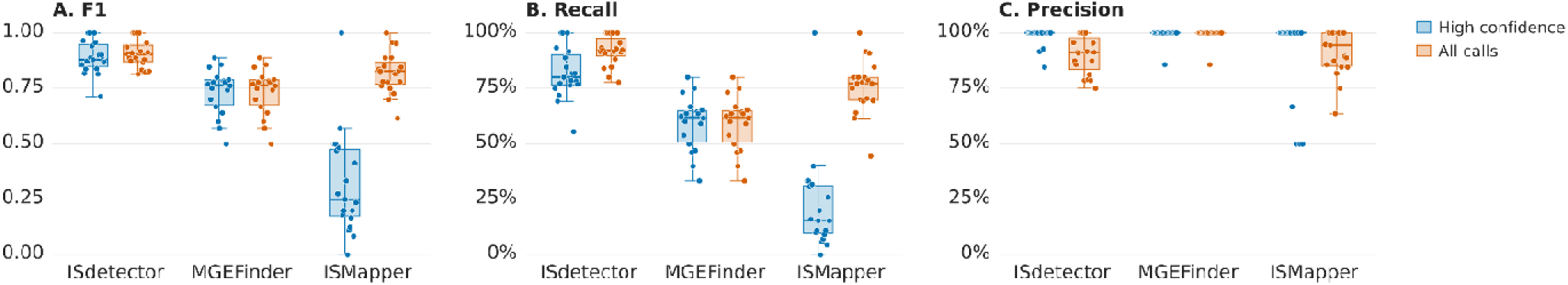
Per-genome performance on 19 real MTB genomes with 50× Illumina reads, using H37Rv as the reference. A. F1 score, B. recall and C. precision for IS6110 at 50-bp tolerance for high-confidence calls (blue) and all calls (orange). Boxes and whiskers are as in Figure 2. MGEFinder panels show 18 genomes because MGEFinder detected no IS6110 insertions in one genome. The ISMapper high-confidence precision panel shows 18 genomes because ISMapper made no high-confidence calls in one genome. Alt text: Plots compare precision, recall, and F1 scores by tool across the real M. tuberculosis benchmark genomes.

**Table 3.**
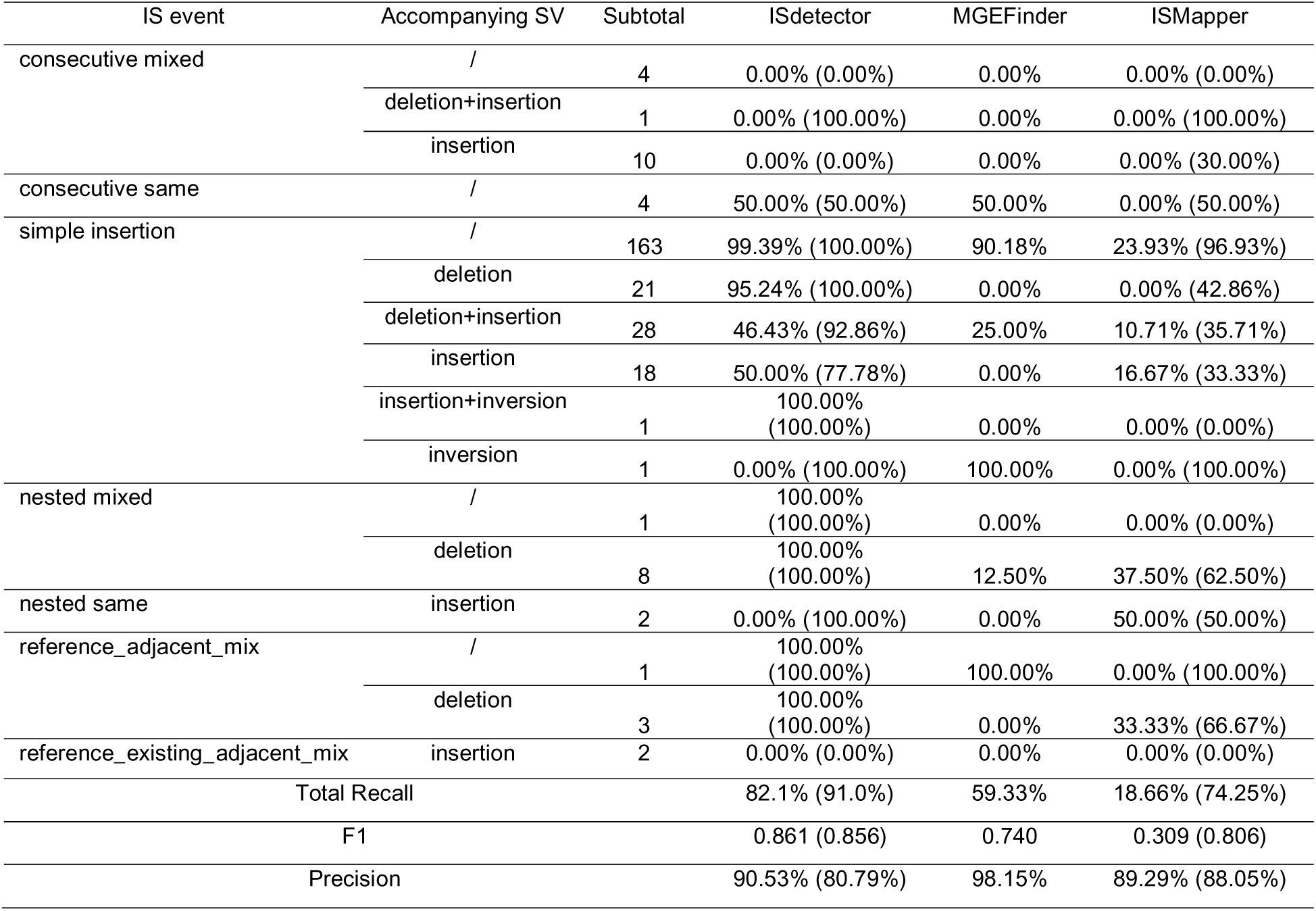
Per-event type recall in real MTB genomes.

| IS event | Accompanying SV | Subtotal | ISdetector | MGEFinder | ISMapper |
|---|---|---|---|---|---|
| consecutive mixed | / | 4 | 0.00% (0.00%) | 0.00% | 0.00% (0.00%) |
|  | deletion+insertion | 1 | 0.00% (100.00%) | 0.00% | 0.00% (100.00%) |
|  | insertion | 10 | 0.00% (0.00%) | 0.00% | 0.00% (30.00%) |
| consecutive same | / | 4 | 50.00% (50.00%) | 50.00% | 0.00% (50.00%) |
| simple insertion | / | 163 | 99.39% (100.00%) | 90.18% | 23.93% (96.93%) |
|  | deletion | 21 | 95.24% (100.00%) | 0.00% | 0.00% (42.86%) |
|  | deletion+insertion | 28 | 46.43% (92.86%) | 25.00% | 10.71% (35.71%) |
|  | insertion | 18 | 50.00% (77.78%) | 0.00% | 16.67% (33.33%) |
|  | insertion+inversion | 1 | 100.00% (100.00%) | 0.00% | 0.00% (0.00%) |
|  | inversion | 1 | 0.00% (100.00%) | 100.00% | 0.00% (100.00%) |
| nested mixed | / | 1 | 100.00% (100.00%) | 0.00% | 0.00% (0.00%) |
|  | deletion | 8 | 100.00% (100.00%) | 12.50% | 37.50% (62.50%) |
| nested same | insertion | 2 | 0.00% (100.00%) | 0.00% | 50.00% (50.00%) |
| reference_adjacent_mix | / | 1 | 100.00% (100.00%) | 100.00% | 0.00% (100.00%) |
|  | deletion | 3 | 100.00% (100.00%) | 0.00% | 33.33% (66.67%) |
| reference_existing_adjacent_mix | insertion | 2 | 0.00% (0.00%) | 0.00% | 0.00% (0.00%) |
| Total Recall |  |  | 82.1% (91.0%) | 59.33% | 18.66% (74.25%) |
| F1 |  |  | 0.861 (0.856) | 0.740 | 0.309 (0.806) |
| Precision |  |  | 90.53% (80.79%) | 98.15% | 89.29% (88.05%) |

There are 105 IS6110 copies accompanied by structural variations or involved in complex insertion events. ISdetector detected 81 of these IS copies, reported SV status of 39 of them, and provided detailed information on 23 deletions, while ISMapper and MGEFinder only detected 41 and 12 of these IS copies, respectively (**Table 3**). Repeated insertions of IS6110 within an IS1547 in the region of NC_000962.3: 888,761-891,469 were observed in seven genomes and the reference genome H37Rv (**Supplementary Figure SF5**). There are four positions of IS6110 insertion within this IS1547 copy in seven samples: 888,866, 888,870, 888,990, and 889,017 (only in H37Rv). ISdetector successfully reported these nested mixed insertions of all eight copies of IS6110 insertions in seven samples. Notably, because 888,990 and 889,017 are separated by only 27 bp, default alignment by minimap2 aligned the IS6110 copy in the sample (NC_000962.3:888,990) to the reference IS6110 copy (NC_000962.3:889,017) even though they were inserted at different reference positions, which introduced two symmetric indels (+28/-27 indels). ISMapper missed both insertions (888,866 and 888,990) in one sample but reported an ambiguous insertion range of 890,388-890,493, and MGEFinder reported no insertions in this region. In another example of complex events, ISdetector successfully reported an IS6110 insertion at 1,986,638 (the closest upstream aligned reference coordinate) accompanied by surrounding deletions (Deletion_1986638_1998622), while ISMapper and MGEFinder both failed for this event (**Supplementary Figure SF6**).

#### 3.3.2 Performance on five *S. sonnei* genomes with simulated short reads

We identified 1,557 copies of five types of ISs in the five complete *S. sonnei* genomes. Because MGEFinder only reports novel insertions, and the *S. sonnei* reference genome already contains hundreds of known IS copies that MGEFinder inherently ignores, the recall of MGEFinder was substantially lower in *S. sonnei*. Therefore, we only compared the performance of ISdetector and ISMapper for real *S. sonnei* genomes. ISdetector showed com arable metrics to ISMapper for both high*-*confidence and all calls (**Figure 5**). No obvious difference in performance was observed among IS types (**Supplementary Table ST7**).

**Figure 5.**
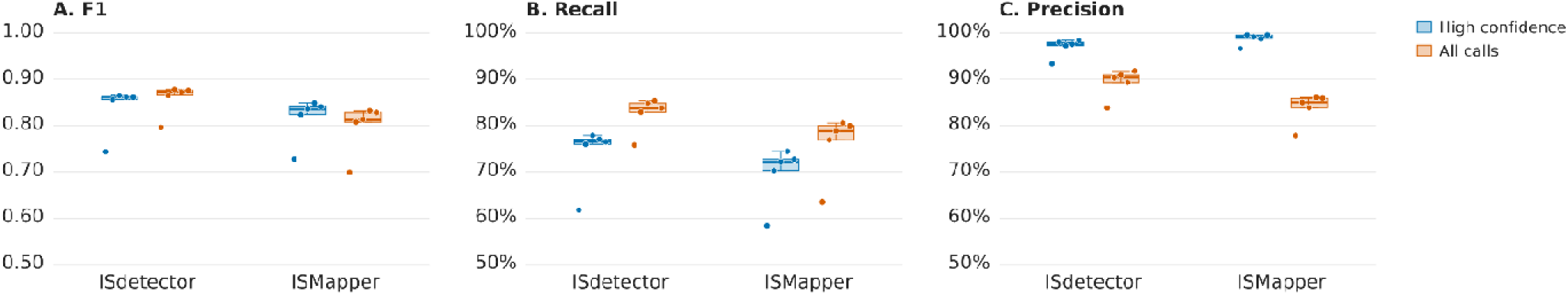
Per-genome performance on five complete *S. sonnei* genomes with simulated 50× short reads, using Ss046 as the reference. A. F1 score, B. Recall and C. Precision for five IS types (IS1A, IS3, IS600, IS2 and IS21) at 50-bp tolerance for high-confidence calls (blue) and all calls (orange). Boxes and whiskers are as in Figure 2. MGEFinder is not shown because it reports only novel insertions and ignores IS copies already present in the reference. Alt text: Plots compare precision, recall, and F1 scores by tool across five S. sonnei genomes.

### 3.4 Comparison of computational efficiency

We compared computational efficiency using four different thread counts (4, 8, 16, and 32) (**Figure 6**). Because MGEFinder requires input in BAM format, its total time included the time to generate BAM files and to run MGEFinder. Increasing thread counts substantially increased RAM usage for all three tools, but the running time benefits varied. MGEFinder running time remained largely unchanged with a slight decrease when increasing from 4 threads to 8 threads, while ISdetector and ISMapper runtimes decreased by approximately one third from 4 threads to 16 threads. Increasing the thread count to 32 slightly increased runtime for both tools. Even with only 4 threads, ISdetector ran substantially faster than either of the other tools at higher thread counts.

**Figure 6.**
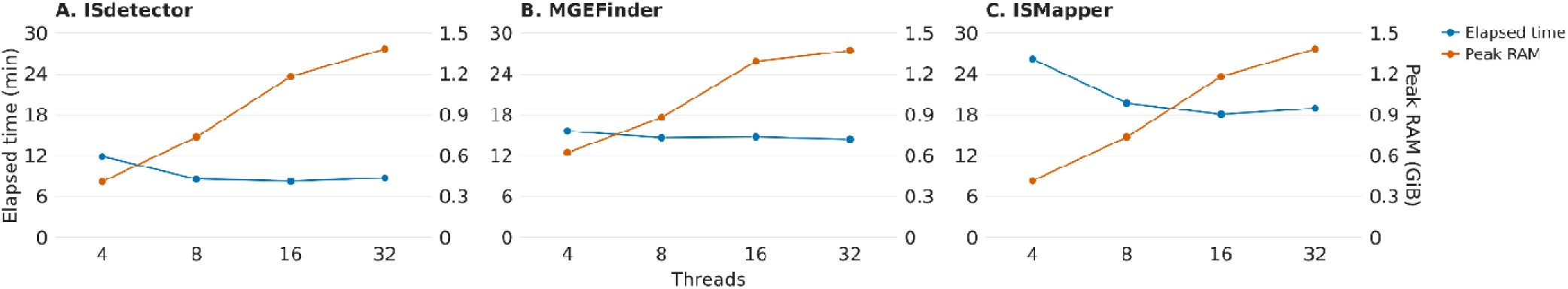
Computational efficiency of A. ISdetector, B. MGEFinder and C. ISMapper at 4, 8, 16 and 32 threads. Elapsed time (left axis, in) is the total for all 19 real MTB genomes; peak RAM usage (right axis, GiB) is the max RAM usage at any time point during the tool running for all 19 MTB genomes. MGEFinder elapsed time includes BAM file generation. Alt text: Plots compare elapsed time and memory use for three tools across four thread counts.

## 4 Discussion

### 4.1 High mapping precision with robust false positive control

ISdetector provides a highly precise and robust solution for mapping the exact positions of ISs and their associated SVs directly from high-throughput short-read sequencing data. The primary achievement of ISdetector lies in its superior performance in both high-IS-burden genomes and high-GC-content genomes for both high-confidence and all calls, achieving high recall across some complex insertion event types, and high accuracy in the predicted insertion positions, whereas existing tools struggled with precision or missed known insertions entirely.

The superior performance of ISdetector is primarily driven by its novel “IS-specific clean reference” and read-clustering methodologies. In genomes with massive IS infestations (such as *S. sonnei*), short sequencing reads can randomly map to multiple identical copies of the repeats scattered across the genome, causing an inflation of false positive results, as observed with ISMapper in real *S. sonnei* genomes, which reported lower precision than ISdetector for all calls. ISdetector solves this “known versus novel” dichotomy by computationally deleting identified IS regions from the original reference to create a synthetic “clean” genome. By mapping reads to this artificial scaffold, both reference-restored and newly inserted IS elements generate uniform breakpoint signals, allowing the pipeline to capture all transposition events using a single, unified logic stream before utilizing a cross-mapping dictionary to translate the coordinates back to the original genome. In high-GC-content genomes with active IS movement such as MTB, ISdetector exhibited much higher recall for both high-confidence and all calls and maintained acceptable precision. In addition, ISdetector offers fine-tuning of parameters, which addresses different needs to increase recall or precision. For example, to reduce FPs, users could increase MIN_PEAK_SUPPORT (e.g. from 1 to 8), which increases precision at the cost of lower recall and F1 score.

### 4.2 Bridging the gap between IS transposition and structural variations

Beyond simply locating insertion sites, ISdetector makes significant progress in bridging the gap between IS transposition and large-scale genomic rearrangements. Transposable elements are notorious for mediating complex structural changes, yet general SV callers often fail to distinguish IS-mediated events from other genomic rearrangements and IS-specific tools often fail to report such IS insertions with other SVs. In our study, ISdetector explicitly identified 81 out of 105 ISs associated with SVs in MTB and reported precise deletion information of 23 of them, far exceeding the capabilities of ISMapper (41/105) and MGEFinder (12/105), which only captured a small fraction of these functionally critical evolutionary events. Especially for complex events involving nearby or nested IS insertions, ISdetector could report highly accurate insertion positions with high recall. By accurately detecting these precise insertion sites and their associated SVs in the coordinates of the annotated reference genome, ISdetector allows researchers to directly link the “mobilome” to phenotypic traits like multidrug resistance and immune evasion, providing a much higher resolution of bacterial evolution than mere presence/absence detection tools.

### 4.3 Limitations and computational trade-offs

Despite its robust performance, a critical analysis of ISdetector reveals inherent limitations tied to the physical constraints of short-read sequencing technologies. The most prominent challenge arises in genomic “hotspots” containing consecutive IS elements. When multiple copies of the same IS type (e.g., an IS6110-IS6110 tandem repeat) are inserted consecutively within a very short distance (such as within 10 bp), the recall of ISdetector for high-confidence calls drops significantly to 24.46–50%. While this is still superior to ISMapper (0–5.18%) and comparable to MGEFinder (23.33–50%), the current algorithm for short reads cannot distinguish multiple insertions at one site and often only reports one or none.

Similarly, ISdetector struggles to identify ISs situated within large insertions, where short sequencing reads (typically 150-250 bp) simply cannot span the breakpoints and uniquely anchor to the flanking reference sequence. For example, in one MTB genome, there is a large insertion of 6,797 bp at position 3,710,380 in region Rv3326-Rv3327, which contains two copies of IS6110 (separated by 3,683 bp). Because the two IS6110 copies are located 408 bp and 263 bp from the start and end of the large insertion, respectively, beyond the span of short reads, neither copy was reported by any of the three tools.

Finally, regarding resource demands, while ISdetector’s multithreading capability saves significant computational time, it comes at the cost of surging RAM usage, meaning ISdetector requires robust computing infrastructure when deployed at a massive population scale. Nevertheless, ISdetector demonstrates that short-read WGS can serve as an efficient and accurate alternative to traditional IS-based typing at the population level. Because IS elements evolve at a rate faster than the accumulation of single nucleotide polymorphisms in highly clonal pathogens, mapping their exact locations is especially insightful for resolving recent transmission chains in local out-breaks of these conservative pathogens such as MTB (Yeh *et al*., 1998) .

In the future, we are planning to integrate long- and short-read data to improve resolution of complex insertions, evaluate graph pangenomes to reduce dependence on a single reference, extend structural variant reporting to inversions and insertions, and assess ISdetector on metagenome-assembled genomes.

## Supporting information

supplementary methods and results

## Acknowledgements

We used AI tools to assist with study design, write scripts for data analysis and visualization, and suggest manuscript revisions for clarity, concision, and alignment with the journal’s guidelines. The authors verified all results and conclusions.

## Data Availability

The 19 MTB complete genomes and their sequencing reads are accessible at China National Center for Bioinformation (PRJCA080033).

## Availability and Implementation

ISdetector is implemented in Python and the archived release are available at https://github.com/carolynzy/ISdetector/releases/tag/v1.1.0.

## Supplementary Information

Supplementary data accompany this article.

---

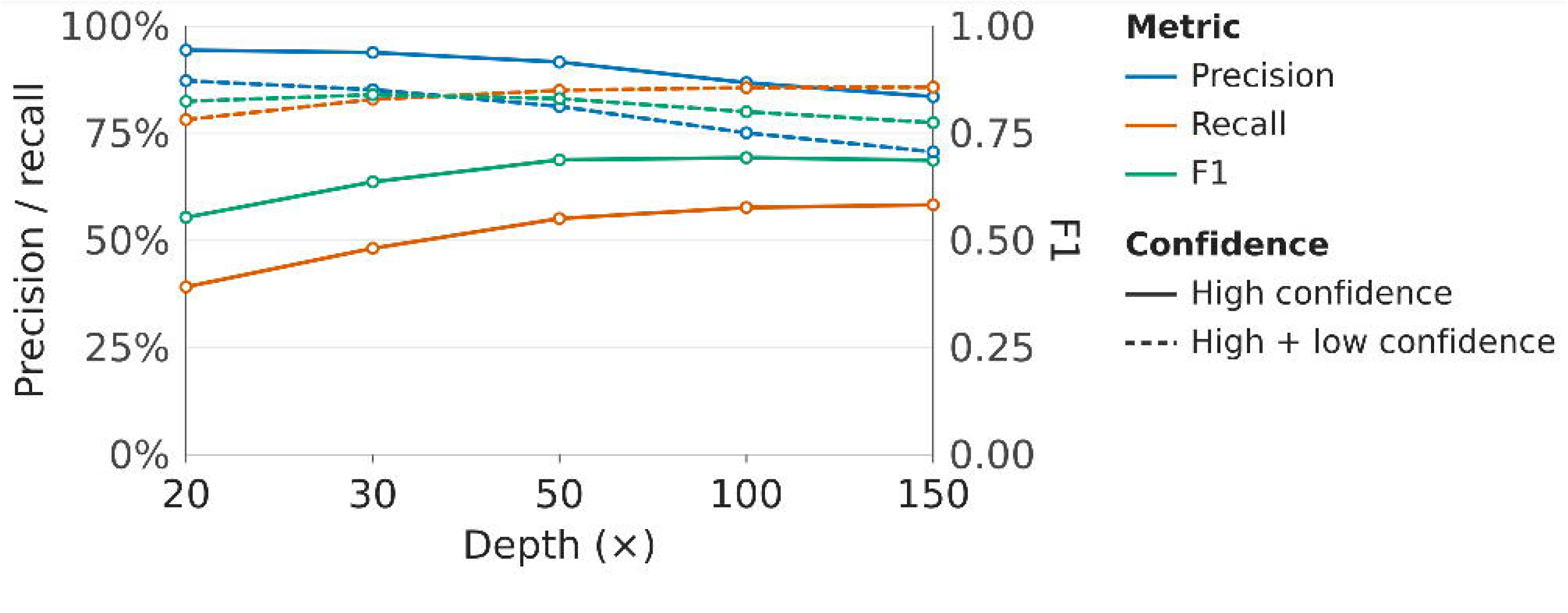

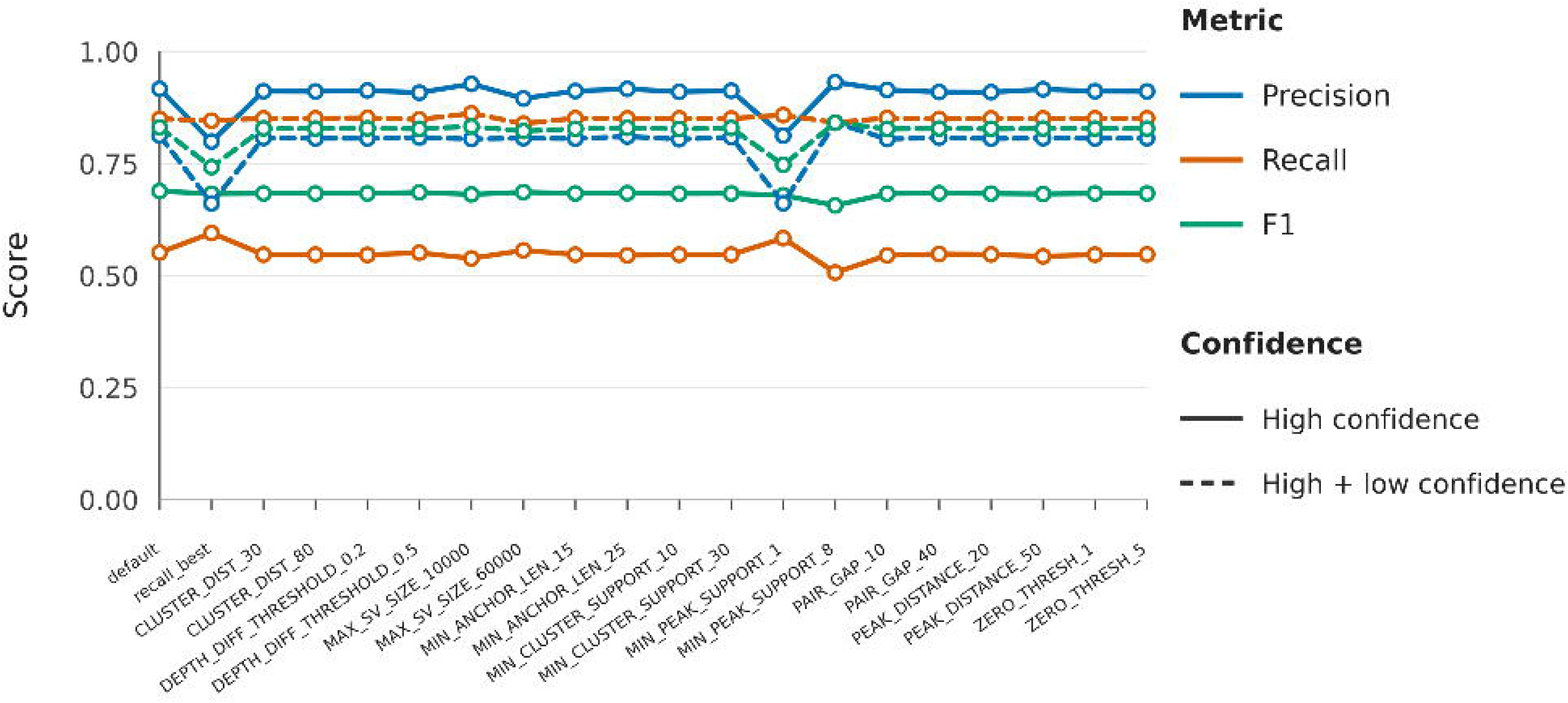

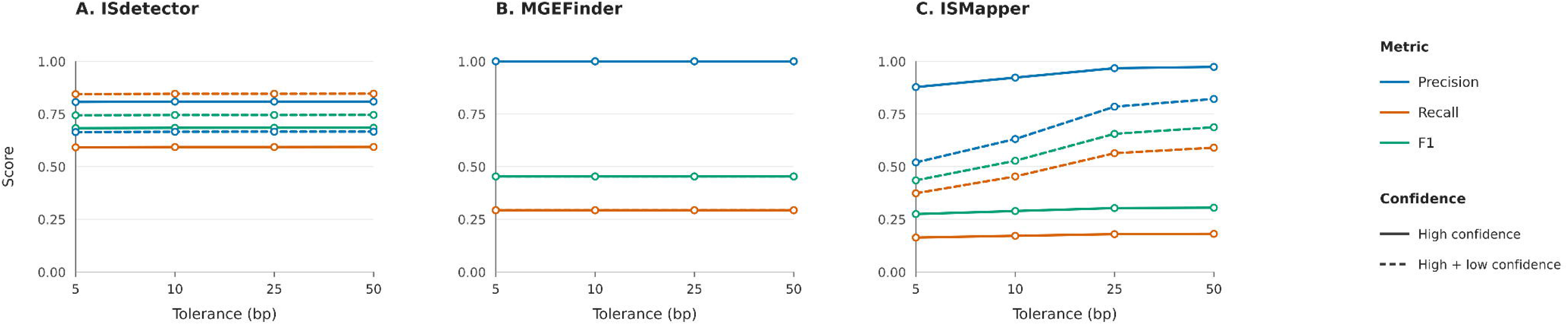

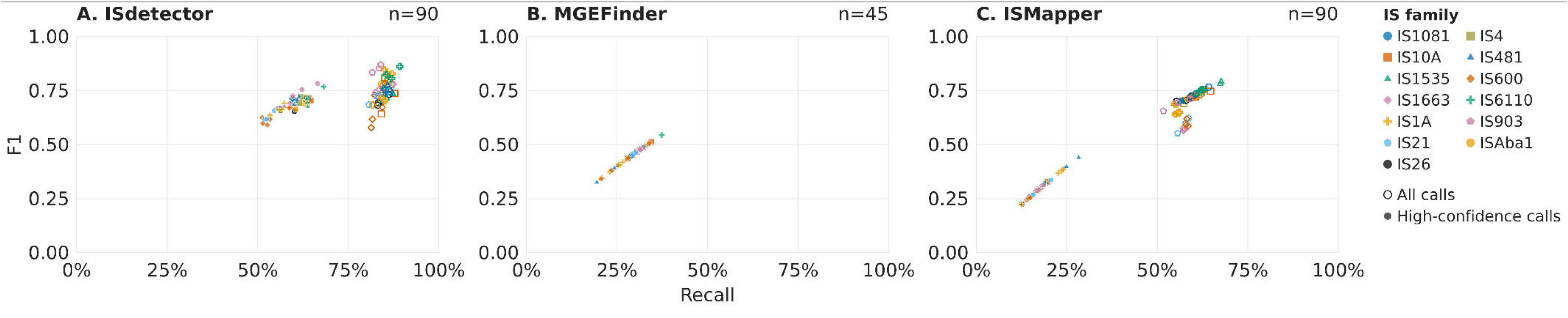

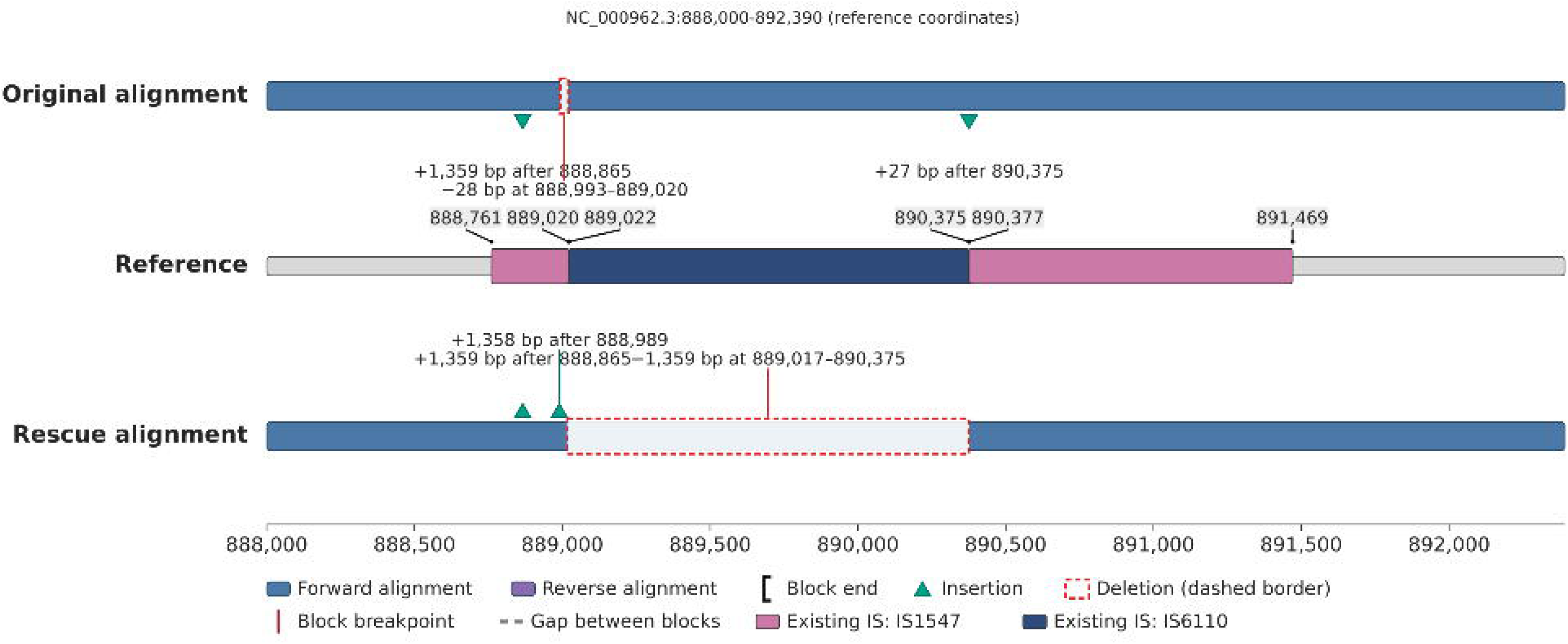

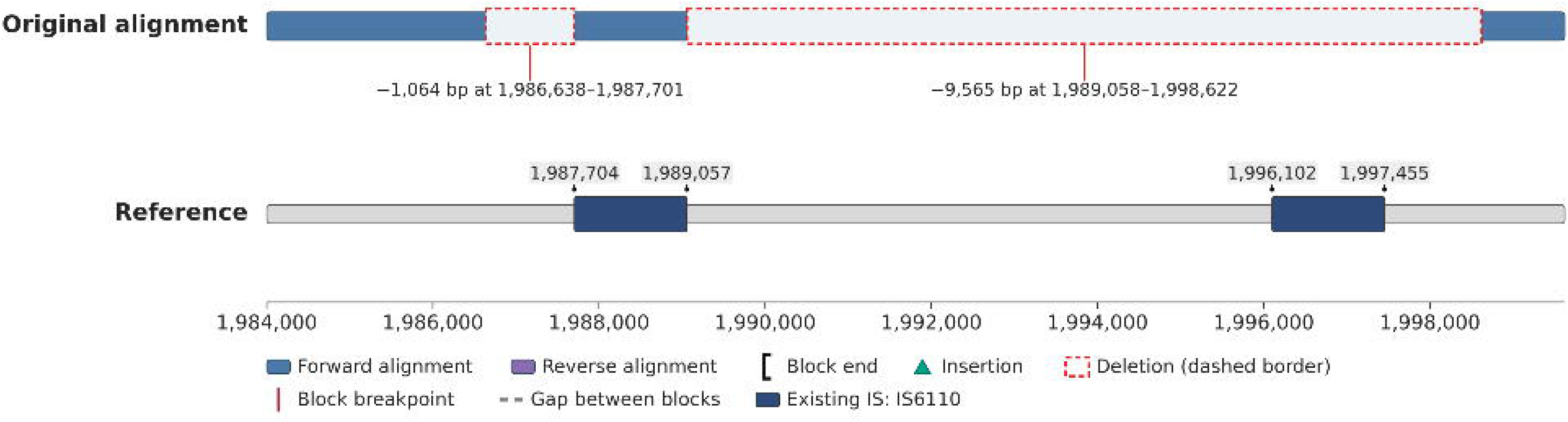

