## supplementary methods and results for "ISdetector: precise mapping of insertion sequences and associated structural variations from short-read sequencing data"

**ISdetector supplementary material**

### Supplementary Methods

#### Modeling factors associated with recall

We fitted separate ridge-penalized logistic regression models for each tool at the level of individual simulated truth events with the same 18 replicate genomes used for batch evaluation. Replicate genomes used for depth sensitivity analysis and profile evaluation are not used in modeling. The binary response indicated whether a tool reported a call that matched the truth event within a 50-bp positional tolerance. False-positive calls were excluded because the analysis focused on recall. For ISdetector and ISMapper, we fitted separate models for all calls and for high-confidence calls.

The full model included 10 predictors: IS–genome type, insertion event type, total IS burden, same-type IS burden, IS length, retained fraction, reference-existing status, nearest-IS distance, local GC content, and flank mappability. Reference genome groups were represented by indicator variables (one per group, six in total). We applied a log (1 + x) transformation to total IS burden, same-type IS burden, and nearest-IS distance, and a natural-log transformation to IS length. These four transformed predictors and local GC content were standardized using the training-fold means and standard deviations. Retained fraction and flank mappability were used without transformation. For categorical predictors, we used the alphabetically first level observed in the training fold as the reference category and represented unseen validation levels with all-zero indicators.

We used nested cross-validation with three outer folds and two inner folds, repeated 10 times with different assignments of genome replicates to folds. Each outer fold held out six of the 18 replicates, one from each reference genome group. The remaining 12 replicates were split in the inner loop to tune the ridge penalty. All events from each replicate stayed together, and preprocessing parameters were estimated within each training fold and applied to its corresponding validation fold.

Within each repetition, we pooled the outer-fold predictions and evaluated log loss, Brier score, ROC AUC, precision–recall AUC, and calibration intercept and slope. To assess the contribution of each predictor block, we refitted a reduced model omitting that block using the same splits and calculated the increase in pooled held-out log loss relative to the full model. We summarized results across the 10 repetitions. All model evaluation used cross-validation within the simulated dataset. No separate independent test set was used.

### Supplementary Results

#### Depth sensitivity

We evaluated sequencing depth using the same set of 15,538 ground-truth IS copies across 12 simulated genomes at each depth. Sensitivity improved most between 20× and 50×. At 50×, high-confidence calls achieved 91.6% precision and an F1 score of 0.688, the second-highest F1 score observed across the depths tested (**Supplement Figure SF1**). Although 100× depth achieved a slightly higher F1 (0.693), it had much lower precision (86.8%).

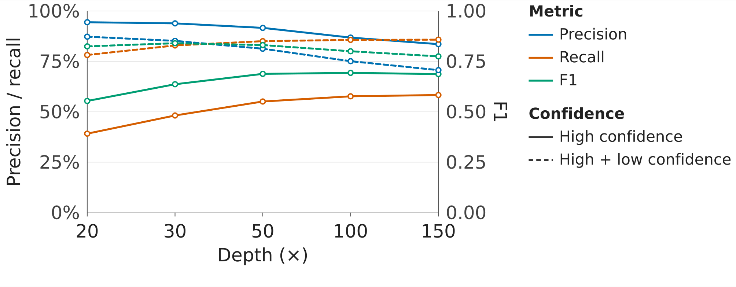

**Supplement Figure SF1.** ISdetector performance across sequencing depths. Precision, recall, and F1 were evaluated at 20×, 30×, 50×, 100×, and 150× using the same 15,538 ground-truth IS copies across 12 simulated genomes. Solid lines show high-confidence calls, and dashed lines show high- and low-confidence calls combined. Precision and recall use the left axis, and F1 uses the right axis.

#### Profile evaluation

We evaluated ISdetector profiles on the same 12 simulated genomes at 50×. High-confidence performance was generally stable across the profiles tested (**Supplement Figure SF2**). The default profile achieved the highest F1 score in the one-factor sweep. Only MIN_PEAK_SUPPORT_1 and MAX_SV_SIZE_60000 improved recall over the default. We combined those two settings into the recall_best profile and evaluated it on the same panel.

Of the 20 profiles tested, recall_best achieved the highest high-confidence recall (59.5%, F1 = 0.68), followed by the best one-factor variant, MIN_PEAK_SUPPORT_1 (58.3%, F1 = 0.68). The default profile achieved 55.1% recall and an F1 score of 0.69.

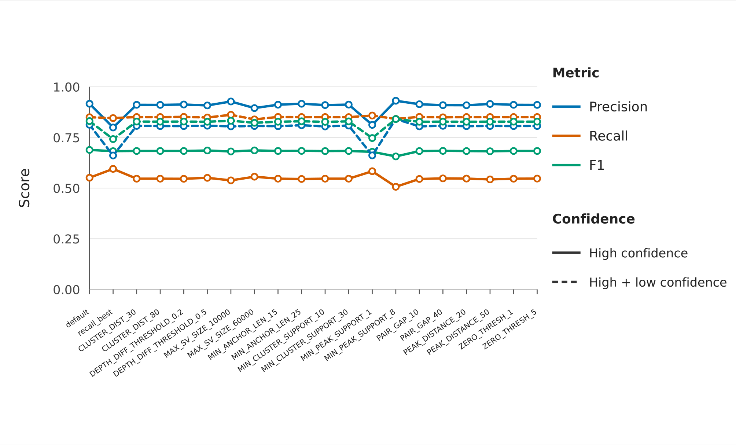

**Supplement Figure SF2. ISdetector performance across parameter profiles.** Precision, recall, and F1 were evaluated for 20 profiles on the same 12 simulated genomes at 50× depth. The recall_best profile combines MIN_PEAK_SUPPORT_1 and MAX_SV_SIZE_60000. Solid lines show high-confidence calls, and dashed lines show high- and low-confidence calls combined.

#### Performance by insertion event type in simulated genomes

Performance differed strongly across insertion types. ISdetector showed the broadest sensitivity across event types (**Supplement Table ST7**). Its high-confidence calls retained strong recall for partial insertions (94.9%), flanking deletions (78.1%), simple insertions (90.6%), nearby insertions (66.3–70.5%), and reference-existing ISs (88.5%). Its main weaknesses were consecutive insertions, nested insertions, reference-adjacent insertions, and events involving flanking insertions or inversions, where high-confidence filtering removed most detections.

For high-confidence calls, ISMapper performed comparably to ISdetector for flanking inversion events and markedly better for flanking insertion events, but it was less sensitive for all other insertion types. MGEFinder had the narrowest event coverage with zero recall for five categories. Its best high-confidence performance was for simple insertions, but recall was very low for the consecutive-insertion, flanking-insertion, and reference-adjacent categories. When all confidence calls were considered, both ISdetector and ISMapper showed great improvement in reacall in all event types, and ISMapper outperformed ISdeteter nested mixed event. However, in other event types, ISdetector maintained superiority, especially in consecutive events, nearby same events, and reference adjacent events.

**Supplement Table ST7. F1, recall, and precision of ISMapper and ISdetector by IS type.**

| Event type | Truth events | ISdetector | MGEFinder | ISMapper |
| --- | --- | --- | --- | --- |
|  |  | High confidence (All confidence) | | |
| consecutive mixed | 3,104 | 22.49% (79.79%) | 0.00% | 0.00% (4.02%) |
| consecutive same | 3,252 | 24.46% (60.37%) | 23.33% | 5.18% (24.19%) |
| flanking deletion | 2,763 | 78.07% (99.39%) | 0.00% | 0.06% (56.56%) |
| flanking insertion | 3,082 | 6.95% (99.22%) | 2.94% | 49.13% (94.61%) |
| flanking inversion | 1,868 | 16.75% (84.76%) | 0.00% | 13.65% (35.87%) |
| nearby mixed | 3,052 | 66.27% (93.40%) | 47.55% | 15.14% (60.18%) |
| nearby same | 3,192 | 70.48% (90.74%) | 45.40% | 0.27% (27.66%) |
| nested mixed | 2,944 | 47.99% (49.70%) | 43.29% | 7.26% (51.22%) |
| nested same | 2,986 | 49.16% (50.63%) | 44.16% | 9.63% (36.68%) |
| partial insertion | 6,296 | 94.93% (99.34%) | 0.00% | 22.39% (90.81%) |
| reference adjacent | 608 | 27.78% (66.26%) | 0.62% | 12.55% (25.72%) |
| reference existing | 1,231 | 88.45% (96.25%) | 0.00% | 65.05% (71.53%) |
| simple insertion | 6,367 | 90.62% (99.28%) | 94.45% | 32.38% (97.83%) |

#### Performance by IS type in simulated genomes

The per-IS results show strong differences among tools (**Supplement Figure SF4**). For both high-confidence calls and all calls, ISdetector achieved the highest recall and F1 score of the three tools for all 15 IS–genome types. When low-confidence and imprecise calls were included, ISMapper showed a greater improvement in recall and F1 score than ISdetector, but its metrics remained lower. MGEFinder achieved 100% precision for every IS–genome type, the highest of the three tools. This reflects its conservative calling behavior rather than superior overall detection.

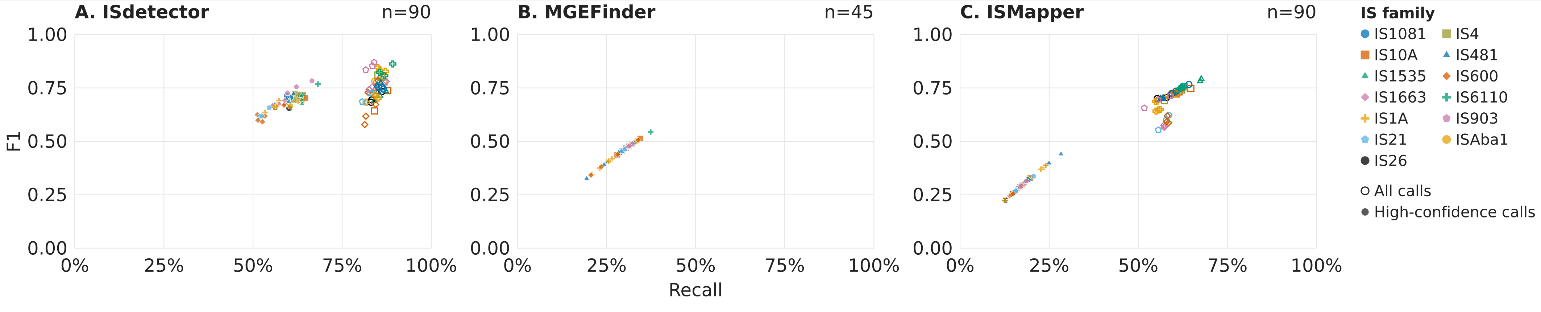

**Supplement Figure SF4. Recall and F1 by IS type in simulated genomes.** Panels A, B, and C show ISdetector, MGEFinder, and ISMapper, respectively. Colors and marker shapes identify IS families. Open markers show all calls, and filled markers show high-confidence calls. Panel labels report the number of plotted observations.
